# Novel model of distal myopathy caused by the myosin rod mutation R1500P disrupts acto-myosin binding

**DOI:** 10.1101/760272

**Authors:** Genevieve C. K. Wilson, Ada Buvoli, Massimo Buvoli, Kathleen C. Woulfe, Lori A. Walker, Leslie A. Leinwand

## Abstract

**Introduction:** More than 400 mutations in β-myosin, a slow myosin motor, can cause both cardiac and skeletal myopathy in humans. A small subset of these mutations, mostly located in the myosin rod, leads to a progressive skeletal muscle disease known as Laing distal myopathy (MPD1). While this disease has previously been studied using a variety of systems, it has never been studied in the mammalian muscle environment. Here, we describe a mouse model for the MPD1-causing mutation R1500P to elucidate disease pathogenesis and to act as a future platform for testing therapeutic interventions.

**Methods:** Because mice have very few slow skeletal muscles compared to humans, we generated mice expressing the β-myosin R1500P mutation or WT β-myosin in fast skeletal muscle fibers and determined the structural and functional consequences of the R1500P mutation.

**Results:** The mutant R1500P myosin affects both muscle histological structure and function and the mice exhibit a number of the histological hallmarks that are often identified in patients with MPD1. Furthermore, R1500P mice show decreased muscle strength and endurance, as well as ultrastructural abnormalities in the SR & t-tubules. Somewhat surprisingly because of its location in the rod, the R1500P mutation weakens acto-myosin binding by affecting cross-bridge detachment rate.

**Conclusions:** While each group of MPD1-causing mutations most likely operates through distinct mechanisms, our model provides new insight into how a mutation in the rod domain impacts muscle structure and function and leads to disease.

## Introduction

Laing distal myopathy (MPD1) is an autosomal dominant disease with variable timing of disease onset that spans from birth into adulthood (1). MPD1 affects skeletal muscle function in a progressive manner: clinically, symptoms begin with weakness in the lower leg anterior compartment that impacts ankle and great toe dorsiflexion (2, 3). Contrary to many other muscle disorders, pathologic findings in muscle biopsies from MPD1 patients are often inconsistent (1).

MPD1-causing mutations have been mapped to the *MYH7* gene that encodes the β-myosin heavy chain, the primary myosin motor expressed in both human heart and in type I, slow skeletal muscle fibers (4). Unexpectedly, only a small number of MPD1 patients also develop a cardiomyopathy, despite higher levels of β-myosin expression in the heart (1, 5–8). MYH7 is a hexameric molecule comprised of a pair of heavy chains and two pairs of non-identical light chains. ∼400 pathogenic mutations causing cardiac and distal myopathies have been identified in both the N-terminal motor domain as well as in the coiled-coil rod region of MYH7 (http://www.hgmd.cf.ac.uk/ac/index.php). The majority of MPD1 mutations are located in the light meromyosin (LMM) domain corresponding to the C-terminal third of the rod that controls assembly of myosin into the thick filaments (6, 9, 10). However, a small number of them are also located in the motor domain (11, 12). MPD1 mutations are primarily codon deletions and missense mutations that introduce a proline residue (6, 9, 10). Both of these genetic defects are predicted to negatively impact the structure of the myosin coiled-coil structure (13). For example, proline residues found in α-helices induce a ∼26-degree kink that could locally unwind the myosin coiled-coil (13).

The biological effects of a subset of MPD1 mutations have been characterized in both non-muscle and muscle cells (14, 15). Muscle cell-based studies have shown that proline rod mutations do not impair incorporation of the mutant myosins into the sarcomere (12) and therefore, do not block formation of the rod coiled-coil structure as originally proposed (9). However, they can trigger myosin cytoplasmic aggregates (12) or cause aberrant myosin packing in thick filaments (16). A progressive dominant hind/fore limb myopathy resembling MPD1 but associated with high frequency of myocardial infarctions has been reported in pigs (17). In this model, sequence analysis revealed an in-frame insertion of two residues (alanine, proline) in MYH7 exon 30; muscle fiber degeneration and regeneration and interstitial fibrosis were also observed. More recently, to characterize the molecular mechanisms of the MPD1-causing mutation L1729del, a *Drosophila melanogaster* model for MPD1 was established (18). By recapitulating some of the morphological muscle defects such as sarcomeric disorganization and myofibril damage observed in patients, this study provided new insight into the pathogenesis of the disease. However, how MDP1 rod mutations act in the mammalian muscle environment remains unclear and understudied. In fact, while numerous genetic mouse models have been developed for studying MYH7 motor domain mutations that cause hypertrophic or dilated cardiomyopathy (19), no mammalian genetic models have yet been reported for examining the effects of myopathy-causing mutations in the rod domain. To fill this research gap, we created the first MPD1 mouse model expressing the R1500P rod mutation identified in a previous study (9). Our data reveal that expression of the mutant myosin affects both muscle histological structure and performance. Transgenic mice show decreased muscle strength and endurance, as well as decreased resistance to fatigue. Remarkably, we also found that the presence of the R1500P rod mutation weakens acto-myosin binding by affecting cross-bridge detachment rate. Since the phenotype of the transgenic mice closely mimics MDP1, we believe that our animal model will be a useful platform for testing and developing future therapeutic interventions for MPD1.

## Results

### Generation and characterization of R1500P mice

To create a mouse model for MPD1, we generated mice expressing the β-myosin R1500P mutation under the control of the well-characterized muscle creatine kinase (MCK) promoter that restricts transgene expression to fast skeletal muscle fibers only. We followed this strategy to circumvent the much lower abundance of slow/Type I fibers in the mouse compared to human (Figure 1A), and therefore increase the opportunity to reveal a muscle phenotype. As previously reported (20), we tracked expression of the transgene by tagging the C-terminal end of the mouse rod domain, which is 99% identical to the human homolog, with a myc-tag epitope. As a control, mice expressing myc-tagged WT β-MyHC were also created (Figure 1A). We obtained 3 transgenic lines for each group, all expressing similar amounts of WT and mutant transgene protein (Figure S1A). Moreover, we found that in tibialis anterior (TA) muscle, their expression levels are ∼35% of total myosin. (Figure 1B). In agreement with the MCK promoter specificity (21), we did not detect transgene expression in any muscles that normally express detectable amounts of Type I/ slow myosin such as soleus and heart (Figure S1B).

**Figure 1.**
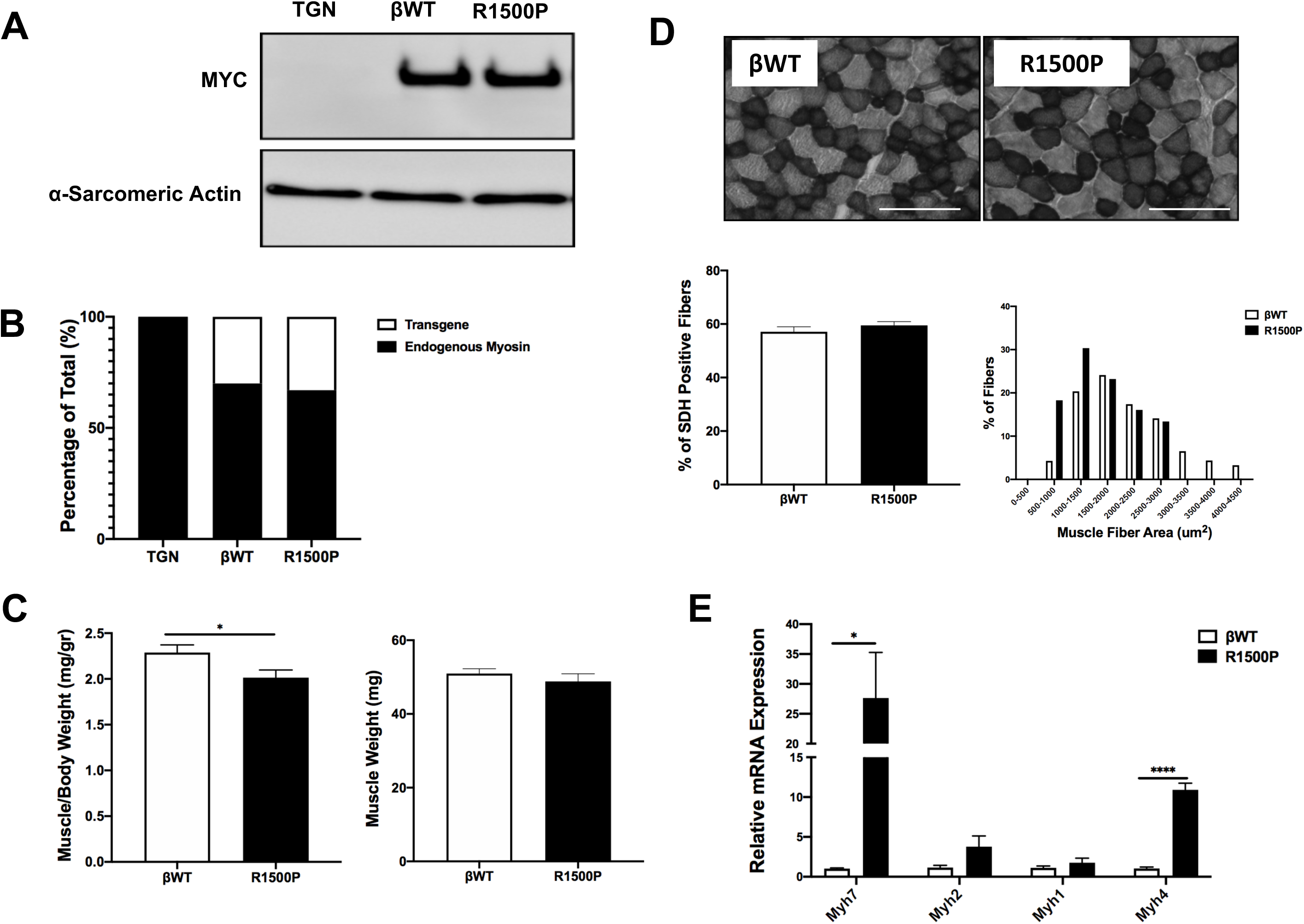
Characterization of R1500P transgenic muscle. (A) Representative western blot analysis performed with tibialis anterior total protein extracts from the indicated genotypes using myc and α-sarcomeric actin (control) antibodies. (B) Quantification of transgenic myosin relative to total myosin in the tibialis anterior muscle. (C) Tibialis anterior muscle weight and muscle/body weight ratio of βWT and R1500P mice (n =20/group). (D) Relative mRNA expression levels of myosin heavy chain isoforms in the tibialis anterior muscle from βWT and R1500P mice (n = 10/group). (E) Representative SDH activity as shown by staining cross-sections of tibialis anterior muscle of βWT and R1500P mice. Quantification of percentage of SDH positive muscle fibers and muscle fiber size are also shown. Data are expressed as mean +/- SEM. *p < 0.05, ****p < 0.0001 by two-tailed unpaired t-test with Welch’s correction.

### Characteristic MPD1 histopathology is present in R1500P muscles

Histological features associated with MPD1 are frequently variable and can include: i) change in muscle fiber size with type I hypotrophy, ii) co-expression of slow and fast myosins, iii) core/minicore structures, iv) mitochondrial abnormalities, and v) muscle necrosis and regeneration (1, 22). Hence, we next determined whether the expression of the R1500P mutant in our mouse model induces some of the pathological phenotypes observed in human muscles. We found that while expression of the R1500P mutant did not change the whole muscle weight of measured fast-type muscles, it significantly decreased the muscle/body weight ratio when compared to the βWT transgenic control (Figure 1C). Histological analysis of WT and R1500P TA muscle enzymatically stained for the mitochondrial enzyme succinate dehydrogenase (SDH) showed no significant difference in the percentage of positive fibers (Figure 1D). However, while the proportion of fast versus slow muscle fibers was unchanged, measurement of fiber cross-sectional area showed that R1500P muscle had a higher proportion of smaller muscle fibers than the βWT control (Figure 1D). Thus, fiber hypotrophy could account for the observed decrease in muscle/body weight. To measure the level of expression of the different myosin heavy chain isoforms in TA muscle, we carried out quantitative real-time PCR (qRT-PCR). We found that RNA for both the slowest and fastest myosin isoforms (Myh7 and Myh4 respectively) were upregulated by R1500P mutant expression, while the RNA level of the intermediate skeletal myosin isoforms Myh2 and Myh1 were not affected (Figure 1E). However, while Myh4 RNA was upregulated, the same was not observed at the protein level. To determine if expression of the R1500P mutant myosin led to sarcomere disorganization, we next complemented these studies by analyzing TA muscle ultrastructure with transmission electron microscopy (TEM). This analysis revealed that the integrity of the major sarcomeric components was not affected. However, ultrastructural changes in the sarcoplasmic reticulum (SR), t-tubules, and mitochondria were observed in the R1500P animals. While WT muscles showed the normal pattern of tightly wound SR networks with accompanying t-tubules triads, R1500P muscles had distended, irregular, and enlarged SR with the t-tubules having a variety of abnormalities ranging from mild to severe dilation of the triad structure (Figure 2A). Furthermore, as confirmed by quantification of mitochondrial DNA content, the number of mitochondria was also decreased (Figure 2B). Taken together, these results demonstrate that a number of the histological hallmarks that are often identified in patients with MPD1 are also present in our transgenic mouse model. Moreover, they provide evidence that the SR structure and/or function could also be altered by the expression of the R1500P mutant myosin.

**Figure 2.**
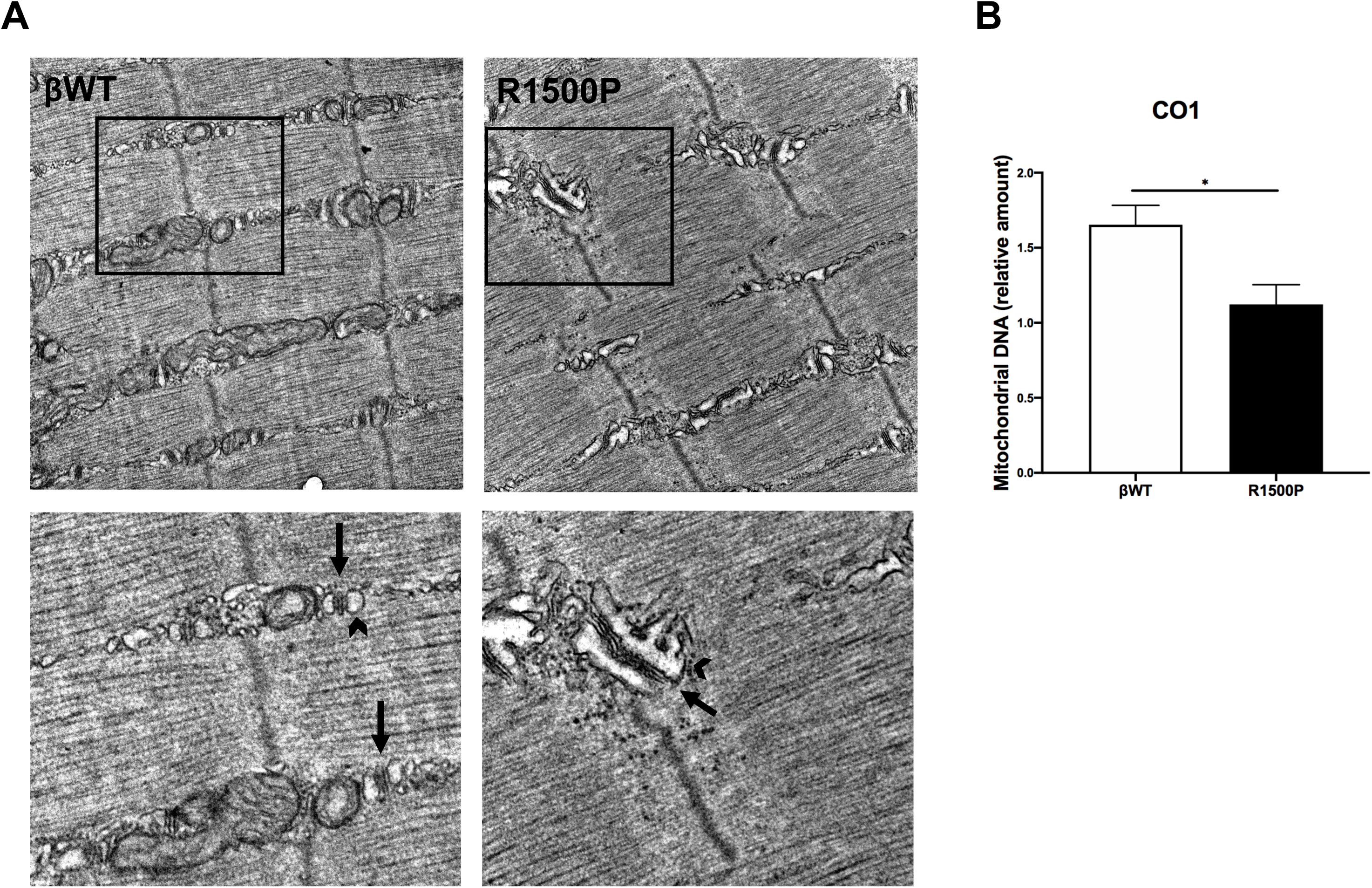
Tibialis anterior muscle shows ultrastructural changes in the SR and t-tubules in the presence of R1500P mutation. (A) In R1500P transgenic mice, t-tubules (black arrows) and sarcoplasmic reticulum (black arrowheads) abnormalities were shown by electron microscopy. βWT TA shows normal t-tubule triads with accompanying organized SR networks. R1500P TA shows dilated t-tubules with distended & enlarged SR. (B) Relative mitochondrial DNA content of CO1 normalized to 18S (n=4/group). Data are expressed as mean +/- SEM. *p < 0.05 by two-tailed unpaired t-test with Welch’s correction.

### The presence of the R1500P mutation activates genes involved in skeletal muscle ER stress and the unfolded protein response (UPR)

Changes and/or disruptions to the ultrastructure of skeletal muscle can cause endoplasmic reticulum (ER) stress, which affects proper ER function by increasing the amount of misfolded/unfolded proteins in the ER lumen (23). As a result, a homeostatic signaling pathway, called Unfolded Protein Response (UPR) is activated. UPR inhibits protein synthesis, increases ER concentration of chaperones, and ultimately, triggers apoptosis (23). Since recent evidence has indicated that UPR is upregulated in a variety of myopathies (26–29), we measured the RNA levels of members of the PERK pathway, which is one of the major UPR transducers and has previously been shown to be activated in muscular dystrophy (26). In TA muscle of R1500P mice, we found significant upregulation of PERK and downstream members of the pathway including ATF4, ATF3, and GADD34 (Figure 3A-D). Other related members of the PERK pathway, such as CHOP were, however, unaffected by the presence of the R1500P mutation (Figure 3E). Thus, in skeletal muscle, heightened activation of genes involved in the UPR pathways may contribute to the development of the MPD1 phenotype.

**Figure 3.**
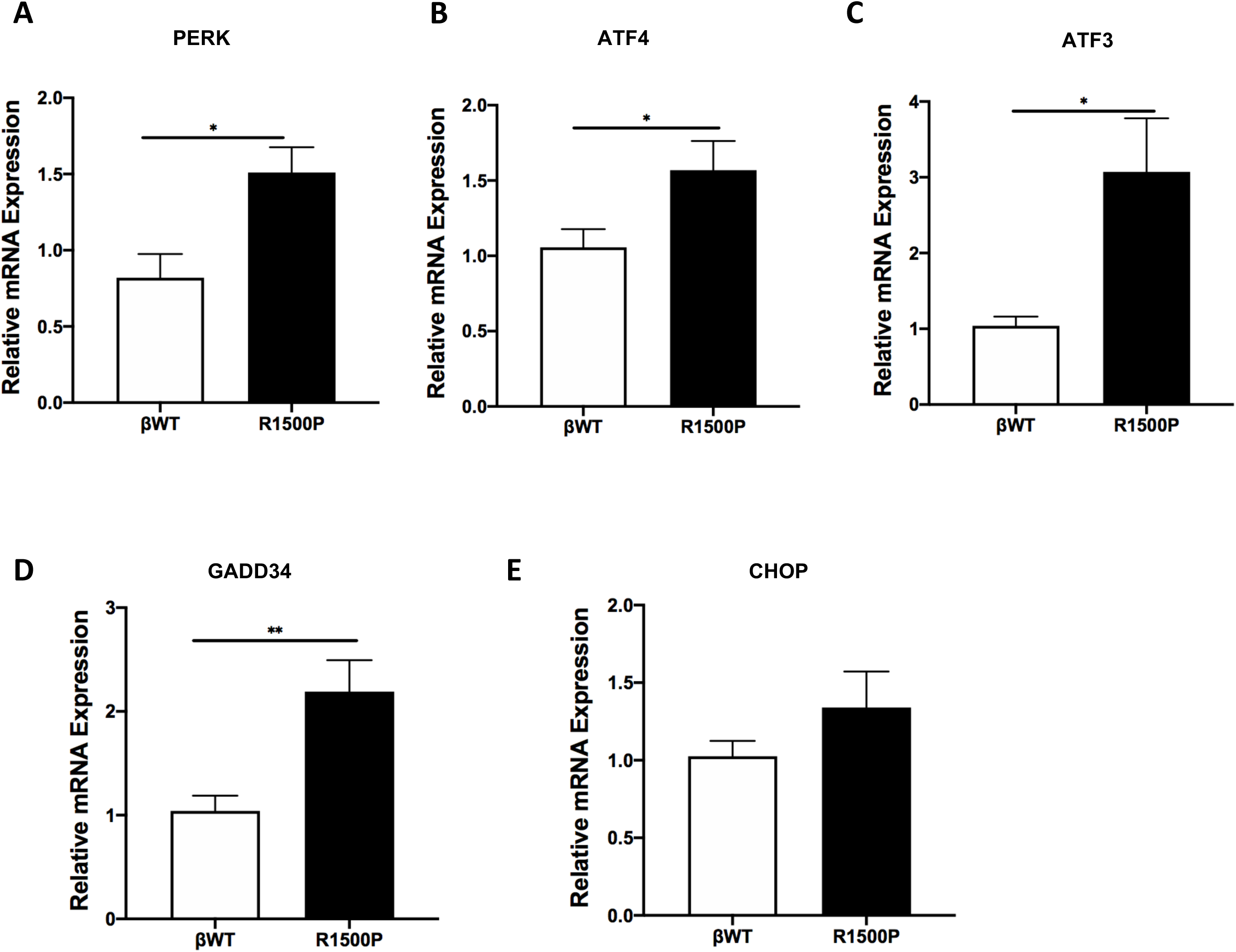
Activation of genes in the unfolded protein response (UPR) in R1500P transgenic mice. (A-E) Relative mRNA expression levels of PERK, ATF4, ATF3, GADD34, and CHOP in the tibialis anterior muscle from βWT and R1500P mice (n = 8/group). Data are expressed as mean +/- SEM. *p < 0.05, **p < 0.01 by two-tailed unpaired t-test with Welch’s correction.

### Muscle fitness and strength are decreased in R1500P mice

To determine whether the R1500P mutation hinders the biochemical mechanics of muscle contraction, we first measured exercise tolerance and fitness in βWT and R1500P mice using a fully automated tracking system to monitor voluntary wheel running. This analysis showed that both the running speed and total running distance over a 28-day period was significantly reduced in 3, 8- and 12-month-old male R1500P mice when compared to their βWT counterparts (Figure 4A). Consistent with these results, muscle strength analyses including a four-limb hanging test and grip strength measurements were impaired with decreased average hang time and a significant reduction in force output in R1500P animals (Figure 4B, 4C). Thus, as seen in MPD1 patients, the expression of the R1500P mutant in our model affects muscle performance and strength.

**Figure 4.**
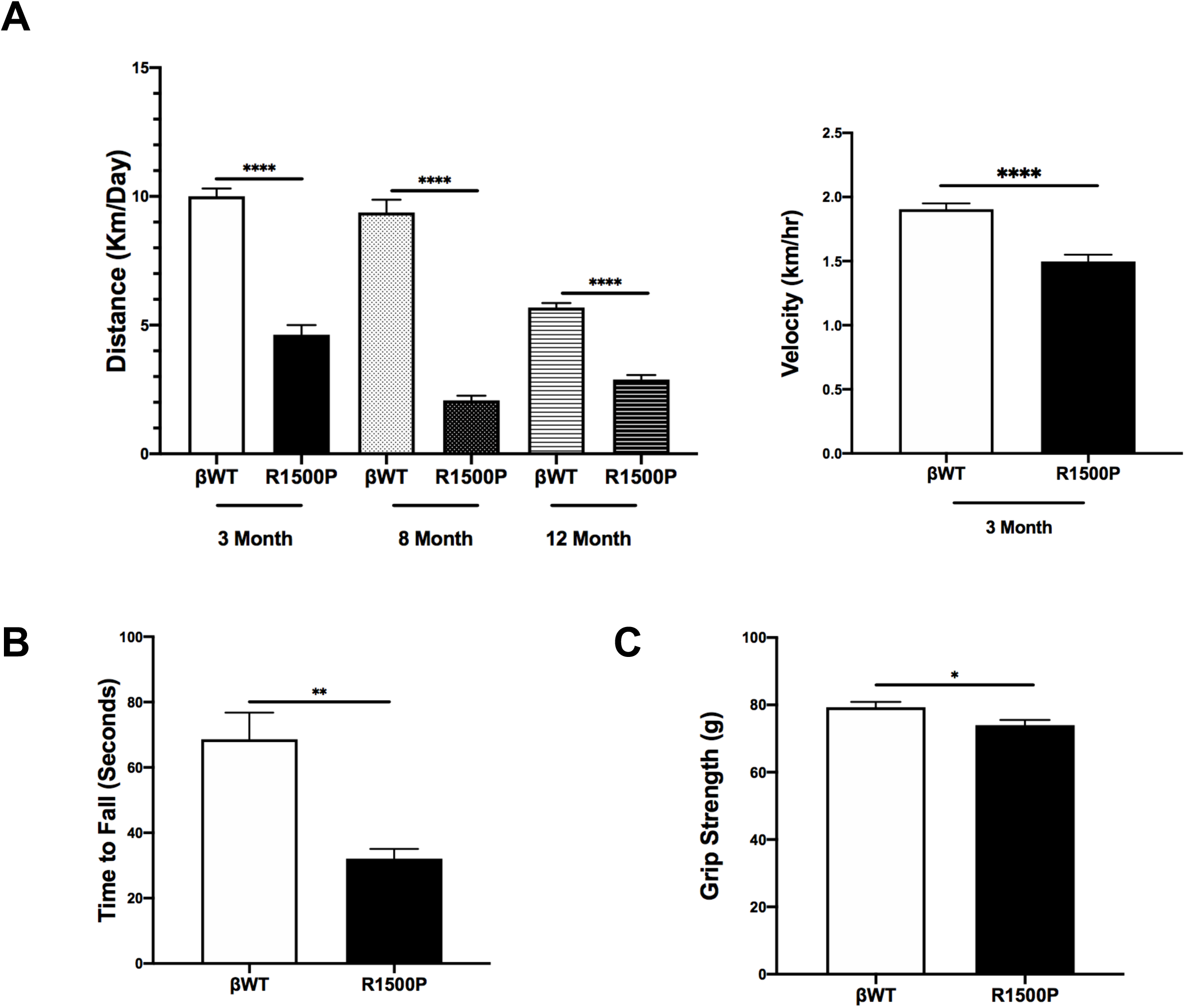
Impaired muscle function in R1500P mutants. (A) Male βWT and R1500P mice were subjected to voluntary wheel running. Bars represent average running distance and speed over 28 days (n = 10-12/group). (B) Four-limb hanging test recording the latency to when the animal falls. Average performance is the average of three trials (n = 6/group). (C) Measurement of forelimb grip strength using computerized grip strength meter (n = 6/group). Data are expressed as mean +/- SEM. *p < 0.05, **p < 0.01, ****p < 0.0001 by two-tailed unpaired t-test with Welch’s correction.

### Ex-vivo contractility assay shows altered R1500P muscle contractility

Since the parameters of muscle contractility, force, fatigability and contractile kinetics can be measured in isolated muscles, we next assessed the *ex-vivo* properties of the extensor digitorum longus (EDL) muscles expressing βWT or R1500P myosins. EDL muscles were used instead of the TA due to the protocol requiring intact muscle-tendon complexes (30). As a control, the properties of the slow-twitch soleus muscle were also analyzed. Measurements of tetanic and twitch force were performed after determining the optimal muscle length. While specific tetanic force was not affected by the presence of the R1500P mutation (Figure 5A), specific twitch force was significantly increased in R1500P EDL muscles while the twitch to tetanus force ratio was significantly decreased (Figure 5B, 5C). In contrast, the activity of WT and R1500P soleus muscles, which do not express the transgene, did not show any functional difference (Figure S2 A-D). Figure 5D shows the force versus frequency relationship for EDL muscles obtained from WT and R1500P EDL muscles. In agreement with the observed decreased twitch/tetanus ratio, the R1500P curve is shifted to the right (downward). Notably, this phenotype, previously characterized as low-frequency fatigue, appears to be linked to altered calcium release (31). We next determined muscle fatigue by challenging the muscles with high frequency stimuli (100 Hz), which induced sustained muscle contraction; as the muscle relaxed over time, we measured the force output as an indicator of fatigue. The measurements obtained showed that the percentage of peak tetanic force dropped more quickly for R1500P muscle (Figure 5E) suggesting that in addition to a drop in force production, the muscles expressing the mutant myosin have decreased resistance to fatigue.

**Figure 5.**
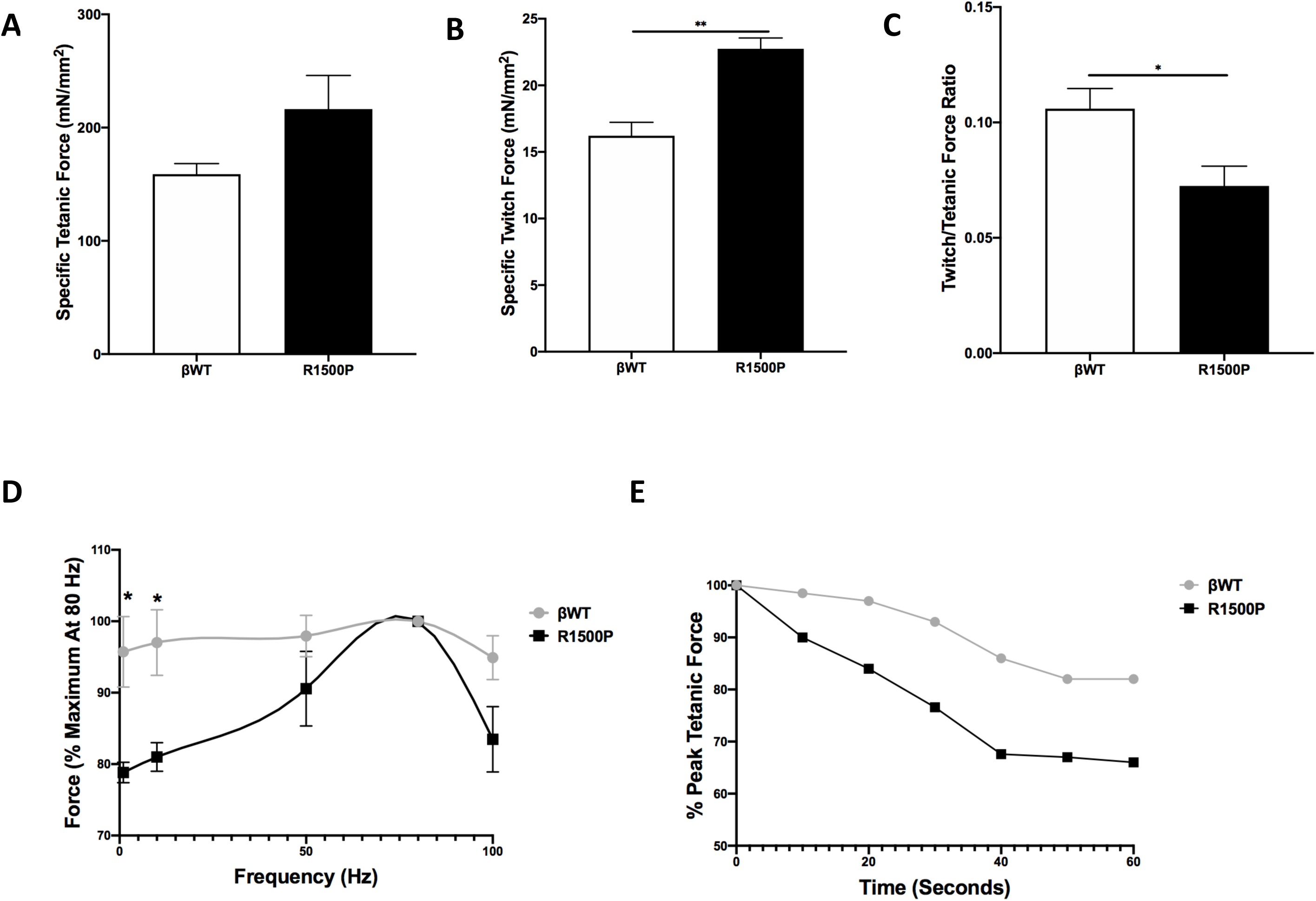
Altered contractility of intact skeletal muscle from R1500P TG mice. (A) Measurement of specific tetanic force in R1500P EDL compared to WT. (B) Measurement of specific twitch force in R1500P EDL compared to WT. (C) Ratio of twitch to tetanic force in βWT and R1500P EDL. (D) Force frequency curves for EDL muscles from βWT and R1500P muscles. Note the right-ward shift of the curve in TG EDL. (E) Percentage of force drop as EDL muscle relaxes after tetanic contraction, measured over 60 seconds (n = 1/group). Data are expressed as mean +/- SEM. *p < 0.05, **p < 0.01 by two-tailed unpaired t-test with Welch’s correction. Unless otherwise noted n = 5/group (βWT), n = 3/group (R1500P).

### The R1500P mutation affects myofibril relaxation

To delve more deeply into the whole animal and whole muscle phenotypes of R1500P mutant mice, we next analyzed force generation and relaxation kinetics of isolated myofibrils from TA muscle. Mounted myofibrils were activated and relaxed by rapidly switching between two flowing solutions of pCa 4.5 and pCa 9.0. After activation, rapid deactivation of myofibrils follows a biphasic state: an initial slow linear decay precedes a faster exponential decay. The rate of slow phase relaxation mirrors cross-bridge detachment rate, whereas the duration of slow phase relaxation depends on Ca^2+^ activation levels. Although we did not detect changes in any mechanical parameters, which include resting, maximal tension, and activation kinetics, k_ACT_ or k_TR._ (Figure S3 A-D), we observed that the rate of slow phase relaxation (k_rel, linear_) was significantly increased in R1500P myofibrils when compared to βWT controls (Figure 6A), whereas the duration of slow phase relaxation (t_rel,linear_) was significantly decreased (Figure 6B). In contrast, the rate of the rapid exponential phase of relaxation (k_rel, exp_) was unchanged (Figure 6C). No difference in activation or relaxation was identified in myofibrils purified from βWT and R1500P soleus muscle (FigureS4 A-E). The faster cross-bridge detachment observed in these experiments conducted on TA indicates that the proline mutation could cause muscle dysfunction muscle by affecting proper acto-myosin binding.

**Figure 6.**
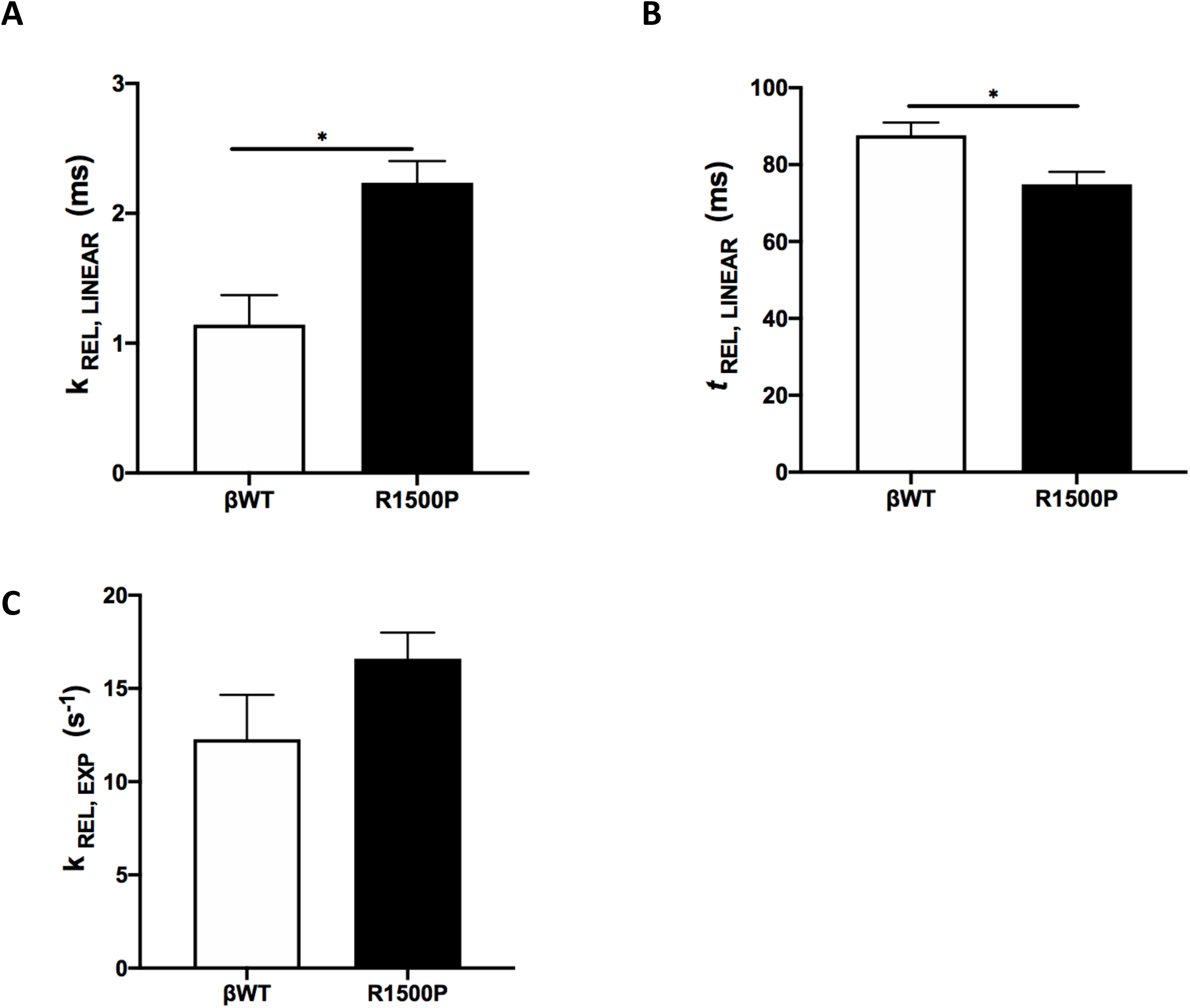
Weaker acto-myosin binding in presence of the R1500P mutation. Measured kinetics of relaxation from myofibrils were as follows: (A) Rate constant of early slow force decline. (B) Duration of early slow force decline. (C) Rate constant of the final exponential phase of force decline. Data are expressed as mean +/- SEM of 6-10 myofibrils/tibialis anterior muscle. *p < 0.05 by two-tailed unpaired t-test with Welch’s correction.

## Discussion

Over 400 mutations in β-myosin have been identified in patients diagnosed with either cardiac or distal skeletal myopathy. A subset of these mutations is known to lead to Laing distal myopathy (MPD1) with mutations in β-myosin being the only known cause of MPD1. However, while this disease has previously been studied in a variety of systems, it is still not understood how these mutations lead to disease – particularly in a mammalian background. In this study, we generated the first mouse model for a mutation in the rod domain of β-myosin.

Patients diagnosed with Laing distal myopathy are known to present with variable muscle pathological changes, which might be due to the position of the mutation within the β-myosin protein – with different mutations resulting in clinical variation. In patients with the R1500P mutation, hypotrophy of slow-type muscle fibers, but variable predominance of type 1 muscle fibers has been noted (9). Histological analyses of muscle from βR1500P muscles shows recapitulation of this phenotype (Figure 1D). While ultrastructural analysis has not previously been performed on patients with the R1500P mutation, the MPD1-causing K1729del has been shown to cause myofibrillar disorganization and mitochondrial abnormalities (1). No sarcomeric disorganization was noted in βR1500P transgenic animals; however, structural abnormalities were noted in the sarcoplasmic reticulum, t-tubules, and mitochondria (Figure 2). Recapitulation of Laing distal histological hallmarks in βR1500P transgenic mice suggests that our model is a useful tool for studying the disease and the underlying pathology.

It was previously proposed that introduction of a proline into the α-helical strands of the myosin rod domain would affect proper assembly of the thick filament due to steric hindrance and the inability to form hydrogen bonds (9, 32). In spite of the predicted effect, previous studies showed that cells transfected with the mutation had organized sarcomeres indicating that the mutant myosin is able to be effectively incorporated into the thick filament – similar to WT (14). However, there are no data about the effect of this mutation on a functioning muscle. Interestingly, Figure 3 shows that mRNAs encoding PERK itself and some downstream effectors were activated implicating ER stress and the unfolded protein response to the R1500P mutant myosin. These results are consistent with structural effects caused by the introduction of a proline residue. While this indicates that the mutant is not as well tolerated as WT myosin and may be contributing to the mechanical phenotypes observed (Figures 5,6), it is also likely that thick filaments containing MPD1 mutants are subject to increased sarcomeric turnover which can be assessed in future experiments.

Our studies showed that the presence of the R1500P mutation in fast-type skeletal muscle led to functional phenotypes, ultrastructural abnormalities in the SR and t-tubules, and contractile deficiencies – on the whole muscle level and at the level of the myofibril. Collectively, the observed phenotypes have all previously been linked to diminished muscle function, effects on cross-bridge detachment rate, and effects on calcium (Ca^2+^) handling. SR & t-tubule enlargement and dis-organization have also been shown to play a role in decreased muscle strength and muscle atrophy (33). Furthermore, force production was shown to be impaired at low frequencies which can lead to insufficient Ca^2+^release preventing full cross-bridge interaction to occur (34). While it remains to be seen how the substitution of a proline affects the structure of the SR/t-tubule network, it would be interesting to determine if there is a change in the amount of Ca^2+^ being released or a change in Ca^2+^ handling which could be leading to the accelerated detachment of myosin cross-bridges from the thin filament.

Here, we have developed the first mouse model for Laing distal myopathy causing mutation R1500P and have shown that these animals display a wide number of phenotypes. While each group of MPD1-causing mutations most likely operates through distinct mechanisms, our model provides new insight into how a mutation in the rod domain may be impacting muscle structure & function and fundamentally leading to disease.

## Materials and Methods

### Animal Care

All animal experiments were performed using protocols approved by University of Colorado Institutional Animal Care and Use Committees (IACUC). Animals were housed under standard conditions in a partial barrier facility and received access to water and chow *ad libitum*. For sample collection, animals were sedated using 1-4% inhaled isoflurane and sacrificed by cervical dislocation. All data shown is from male mice.

### Western Blotting

Protein lysates were prepared by homogenizing hindlimb muscle tissue in myosin extraction buffer (0.3M NaCl, 0.1M NaH_2_PO_4_, 0.05M Na_2_HPO_4_, 0.001M MgCl_2_.6H_2_O, 0.01M EDTA) following standard procedures. The antibodies used were against Myc-Tag (9B11) (1:10000, Cell Signaling Technology, #2276), F59 (1:2000), and α-sarcomeric actin (1:2000). All blots were imaged using the ImageQuant LAS 4000 (GE Healthcare Bio-Sciences, Pittsburg, PA) system and analyzed with the ImageQuant software and/or with ImageJ.

### RNA Isolation & Quantification

Total RNA was purified from hindlimb muscles using TRI Reagent (Ambion) according to manufacturer’s protocol. cDNA was synthesized using SuperScript III reverse transcriptase (Invitrogen) and random hexamer primers. Gene expression was determined by qRT-PCR using SYBR Green dye (Invitrogen) and gene specific primer sets. All genes were normalized to 18S expression. Data were collected and analyzed using Bio-Rad CFX Real-Time PCR system.

### Histology (Succinate Dehydrogenase Staining)

Tibialis anterior muscles were snap frozen in isopentane/liquid N_2_, cryo-sectioned, and stained for enzymatic activities using standard procedures. The stained fibers were counted and their percentage of total number of fibers was calculated (150–200 total fibers/image, 3 images/mouse, 2 mice/genotype). Cross sectional area was determined using ImageJ.

### Transmission Electron Microscopy

Skeletal muscle was dissected and immersed in 2.5% glutaraldehyde and 2% paraformaldehyde in 0.1 M cacodylate buffer at pH 7.4 for a minimum of 24 hours at 4°C. For processing, the tissue was rinsed in 100 mM cacodylate buffer and then immersed in 1% osmium and 1.5% potassium ferrocyanide for 15 min. Next, the tissue was rinsed five times in cacodylate buffer, immersed in 1% osmium for 1 hour, and then rinsed again five times for 2 min each in cacodylate buffer and two times briefly in water. The tissue was stained en bloc with 2% uranyl acetate for 1 hour before it was transferred to graded ethanols (50, 70, 90, and 100%) for 15 minutes each. Finally, the tissue was transferred through propylene oxide at room temperature and then embedded in LX112 and cured for 48 h at 60°C in an oven. Ultra-thin sections (55 nm) were cut on a Reichert Ultracut S from a small trapezoid positioned over the tissue and were picked up on Formvar-coated slot grids or copper mesh grids (EMS). Sections were imaged on a FEI Tecnai G2 transmission electron microscope (Hillsboro, OR) with an AMT digital camera (Woburn, MA).

### Functional Phenotypic Measurements

#### Voluntary Wheel Running

Male mice were subjected to voluntary wheel running for a period of 28 days at the age of 3 months, 8 months, and 12 months. Mice were housed individually in a large cage with a running wheel. Exercise time, velocity, and distance were recorded daily for each animal.

#### Grip Strength

Forelimb grip strength was measured with a grip strength meter. The mice were first acclimated to the apparatus for approximately 5 min. Individual mice were then allowed to grab the bar while being held from the tip of their tail. The mouse was gently pulled away from the grip bar. When the mouse could no longer grasp the bar, the reading was recorded. Protocol was repeated five times with at least 30 sec rest between trials. The highest three values were averaged to obtain the absolute grip strength.

#### Four Limb Hanging Test

Male mice were placed in the center of a wire mesh screen, a timer was started and the screen was rotated to an inverted position with the mouse’s head declining first. The screen was held above a padded surface. Either the time when the mouse falls was noted or the mouse was removed when the criterion time of 60 sec was reached.

### Ex-Vivo Contractility Assay

Mice were euthanized according to NIH guidelines and IACUC institutional animal protocols. Extensor digitorum longus (EDL) was carefully dissected in total from the ligamentary attachment at the lateral condyle of the tibia to the insertion region. The muscle was transferred to a dish containing ice-cold isotonic physiologic salt solution (Tyrode’s buffer (mM): NaCl 118, KCl 4, MgSO_4_ 1.2, NaHCO_3_ 25, NaH_2_PO_4_ 1.2, glucose 10 and CaCl_2_ 2.5) bubbled with 95% O2/5% CO2 to maintain a pH of 7.4. The soleus was identified after removing the gastrocnemius muscle and was removed by cutting the ligaments connecting to the proximal half of the posterior tibia to the insertion, where the calcaneal tendon was cut and the muscle was placed into ice-cold Tyrode’s buffer. Muscles were mounted vertically in individual tissue bath chambers and maintained at 37° C. Muscles were stretched and optimal length was set for each muscle. Stimulatory trains of varying frequency (1-100 Hz) were used to generate force-frequency curves. Tetanic force was achieved in all muscles using 100 Hz.

### Myofibril Isolation

Myofibrils were isolated from flash frozen soleus and tibialis anterior as described (35, 36). A small section of muscle was cut into thin slices and bathed in 0.05% Triton X-100 in Linke’s solution (132mM NaCl, 5mM KCl, 1mM MgCl_2_, 10mM Tris, 5mM EGTA, 1mM NaN_3_, pH 7.1) with protease inhibitor cocktail (10 µM leupeptin, 5 µM pepstatin, 200 µM phenyl-methylsuphonylfluoride, 10 µM E64, 500 µM NaN_3_, 2 mM dithioerythritol) overnight at 4°C overnight. Skinned tissue was washed three times in rigor solution (50mM Tris, 100mM KCl, 2mM MgCl_2_, 1mM EGTA, pH 7.0) and resuspended in bath solution with protease inhibitors (pCa 9.0; 100mM Na_2_EGTA; 1M potassium propionate; 100mM Na_2_SO_4_; 1M MOPS; 1M MgCl_2_; 6.7mM ATP; and 1mM creatine phosphate; pH 7.0) and homogenized at medium speed for 10 seconds three times.

### Myofibril Mechanics

Myofibrils were isolated and mechanical parameters were measured as described (36– 38). Myofibrils were placed on a glass coverslip in relaxing solution at 15°C and then a small bundle of myofibrils was mounted on two microtools. One microtool was attached to a motor that produces rapid length changes (Mad City Labs) and the second microtool was a calibrated cantilevered force probe (5.8 µm/µN; frequency response 2-5 KHz). Myofibril length was set at 5-10% above slack length and average sarcomere length and myofibril diameter were measured using ImageJ. Mounted myofibrils were activated and relaxed by rapidly translating the interface between two flowing streams of solutions of pCa 4.5 and pCa 9.0 (38, 39). Data was collected and analyzed using customized LabView software. Measured mechanical and kinetic parameters were defined as follows: resting tension (mN/mm^2^) – myofibril basal tension in fully relaxing condition; maximal tension (mN/mm^2^) – maximal tension generated at full calcium activation (pCa 4.5); the rate constant of tension development following maximal calcium activation (*k*_ACT_); the rate constant of tension redevelopment following a release-restretch applied to the activated myofibril (*k*_TR_) (40); rate constant of early slow force decline (*k*_REL, LIN_) - the slope of the linear regression normalized to the amplitude of relaxation transient, duration of early slow force decline - measured from onset of solution change to the beginning of the exponential force decay, the rate constant of the final exponential phase of force decline (*k*_REL, EXP_).

### Data & Statistical Analyses

Data are presented as mean ± SEM. Differences between groups were evaluated for statistical significance using Student’s two-tailed t test (two groups) or one-way ANOVA (more than two groups) followed by Tukey’s post-hoc test for pairwise comparisons. *P* values less than 0.05 were considered significant unless otherwise noted.

## Supplementary Figure Legends

**Figure S1. Establishment of βWT and R1500P transgenic mouse lines.**

(A) Representative western blot analysis of transgene expression from established lines performed with tibialis anterior total protein extracts from the indicated genotypes using myc and α-sarcomeric actin (control) antibodies.

(B) Western blot analysis of transgene expression in different tissue types performed with total protein extracts from the indicated tissues using myc and α-sarcomeric actin (control) antibodies.

**Figure S2. No change in contractility of intact soleus muscle from TG mice.**

(A) Measurement of specific tetanic force in R1500P soleus compared to βWT.

(B) Measurement of specific twitch force in R1500P soleus compared to βWT.

(C) Ratio of twitch to tetanic force in βWT and R1500P EDL.

(D) Force frequency curves for soleus muscles from βWT and R1500P muscles.

Data are expressed as mean +/- SEM.

**Figure S3. R1500P mutation does not affect myofibril contractility activation.**

Measured activation kinetics from myofibrils were as follows:

(A) The rate constant of tension development following maximal calcium activation.

(B) The rate constant of tension redevelopment following a release-restretch applied to the activated myofibril.

(C) Myofibril basal tension in fully relaxing condition.

(D) Maximal tension generated at full calcium activation (pCa 4.5).

Data are expressed as mean +/- SEM of 6-10 myofibrils/tibialis anterior muscle.

**Figure S4. No effect on activation or relaxation kinetics in soleus muscle.**

Measured kinetics of relaxation and activation from myofibrils were as follows:

(A) Rate constant of early slow force decline.

(B) Duration of early slow force decline.

(C) Rate constant of the final exponential phase of force decline.

(D) The rate constant of tension development following maximal calcium activation.

(E) The rate constant of tension redevelopment following a release-restretch applied to the activated myofibril.

Data are expressed as mean +/- SEM of 6-10 myofibrils/soleus muscle.

