## Supplementary figures and images for "Novel model of distal myopathy caused by the myosin rod mutation R1500P disrupts acto-myosin binding"

### Supplementary Figure 1

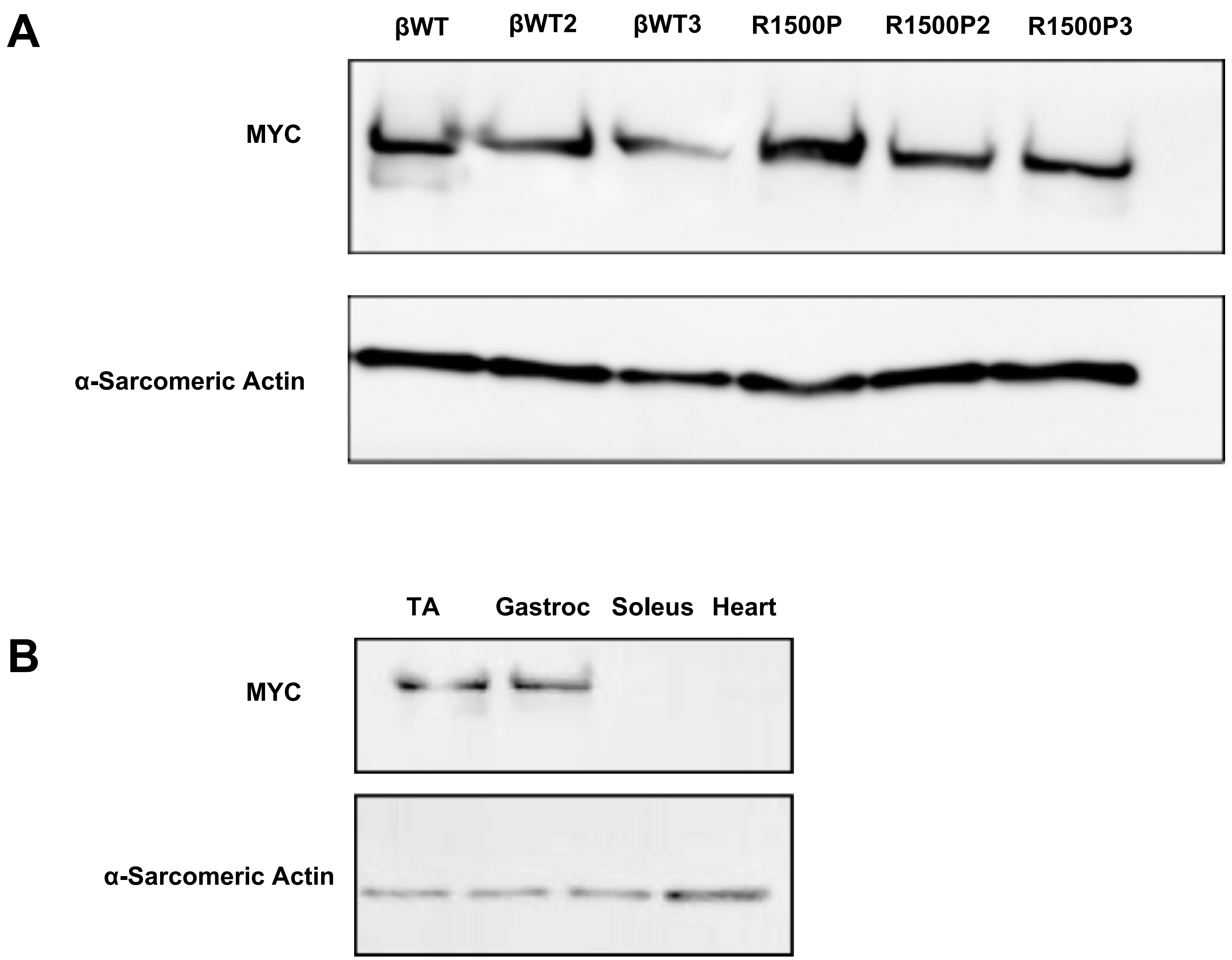

### Supplementary Figure 2

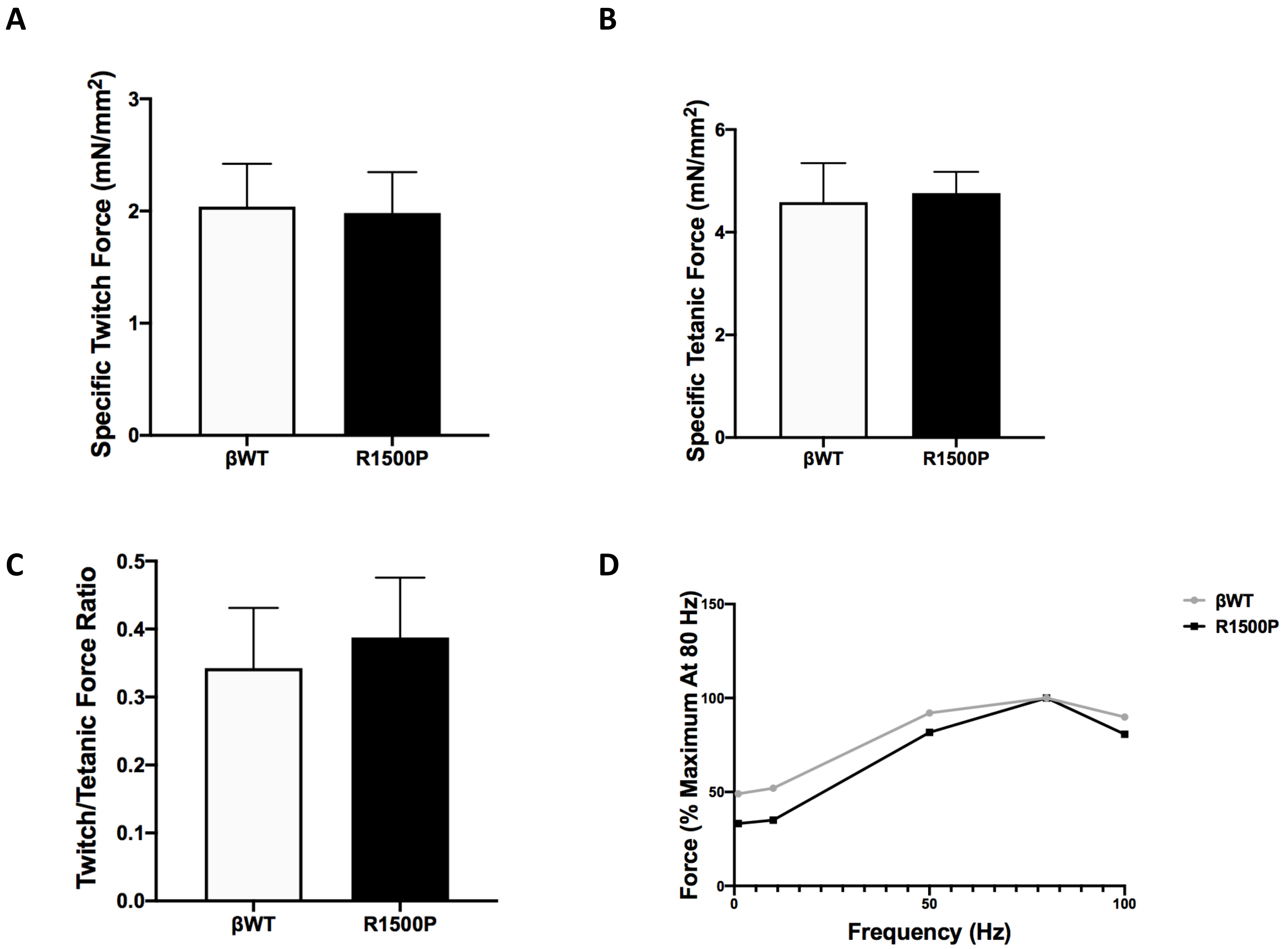

### Supplementary Figure 3

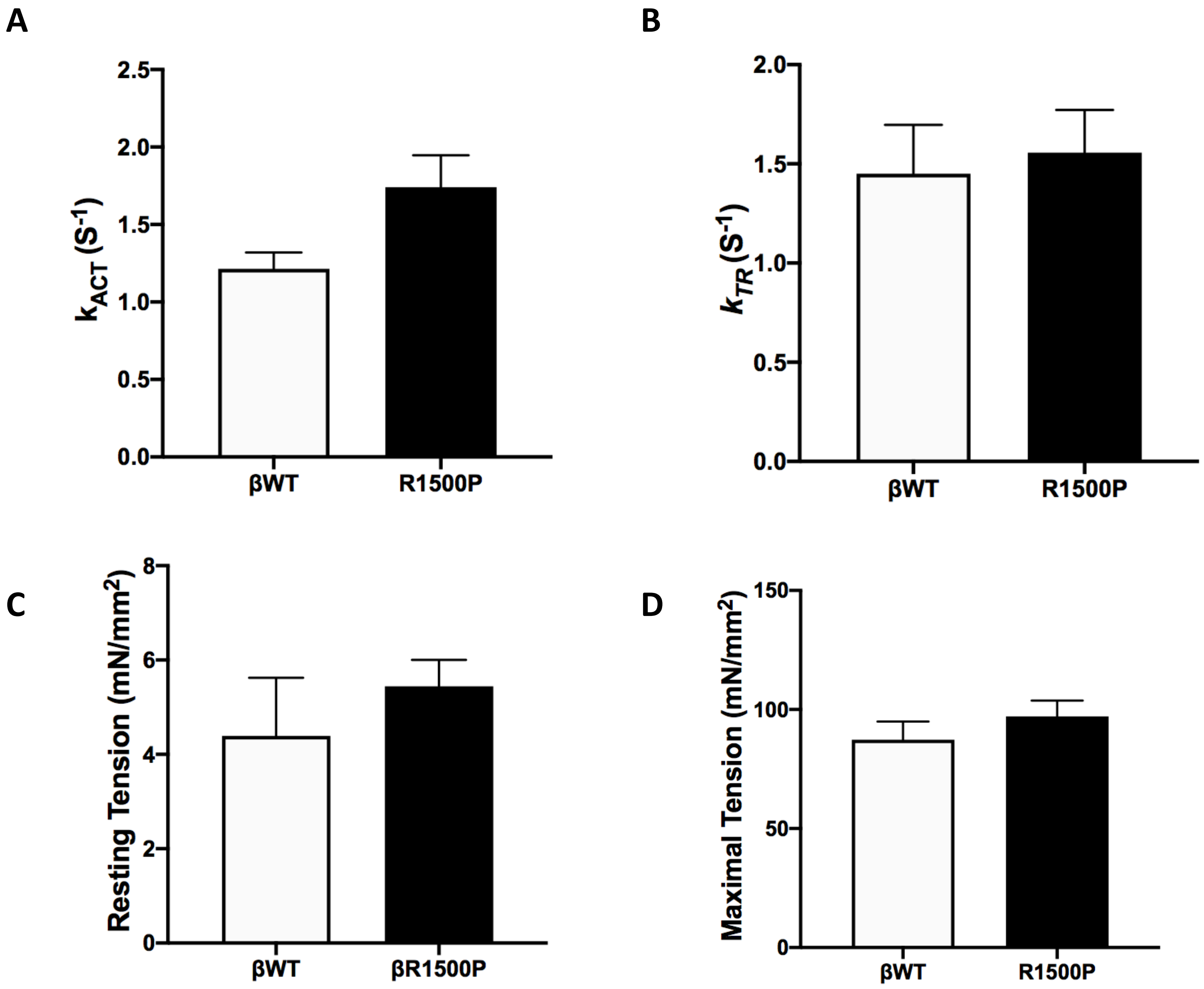

### Supplementary Figure 4

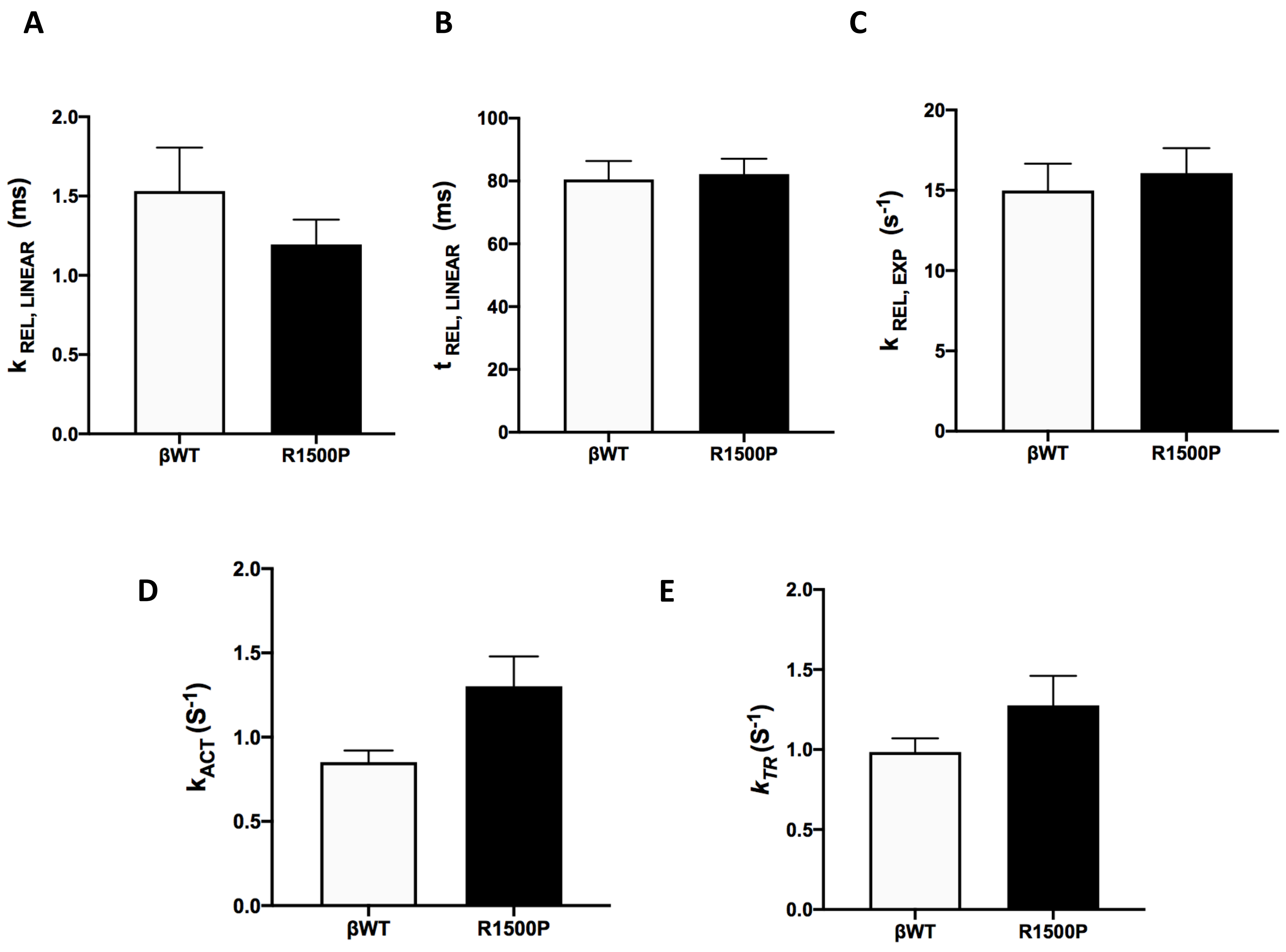
